# Deep learning molecular interaction motifs from Receptor structure alone

**DOI:** 10.1101/2025.01.13.632880

**Authors:** Seeun Kim, Simaek Oh, Hyeonuk Woo, Jiho Sim, Chaok Seok, Hahnbeom Park

## Abstract

Interactions of proteins with other molecules are often mediated by a set of critical binding motifs on their surfaces. The majority of traditional binder design relied on motifs borrowed from known binder molecules, which highly restricted its applicability to novel targets or to new binding sites. In this work, we present a deep learning network MotifGen that predicts potential binder motifs directly from receptor structures without any further supporting information. MotifGen generates motif profiles at the receptor surface for 14 types of functional groups or 6 chemical interaction classes. These profiles are not only highly human interpretable, but also provide pre-trained embedding inputs for versatile few-shot binder design applications. We demonstrate MotifGen’s effectiveness through its application to peptide binder design and small-molecule binding site prediction, where it either surpassed existing methods or added significant value when integrated together. We expect our motif-centric binder design strategy can facilitate discovering novel binders for challenging receptor targets.

## Introduction

Most biological pathways and disease therapeutics are mediated by biomolecular interactions. Despite the diversity of their functions and structures, there are a few key atomic interactions such as hydrogen bonding, hydrophobic interactions, and pi-pi stacking, which largely contributes to the overall molecular interactions. For instance, protein-nucleic acid interactions are strongly mediated by several key hydrogen bonds; many small-molecules bind to nucleic acids through stacking to base(s); protein-protein interactions are known to be mainly driven by the packing of several hydrophobic residues exposed to solvent at unbound state^1,2,3^. Each partner involved in these key interactions contributing to the binding event, often composed by a chemical functional group, is referred to as a “binding motif” in this work.

Given their importance in mediating interactions, various *in silico* methods have been developed to predict such binding motifs. A group of approaches are template-based methods, which leverage prior knowledge of known binders. Depending on the binder molecule type and the way how it is processed, this common direction is called in many different names, such as “ligand-based drug discovery” for small-molecules^4^, “peptidomimetics” for peptides or small proteins^5^, or “motif grafting” for proteins or enzymes^6^.

In contrast to these template-based approaches, identifying novel binding motifs is an essential part in real-world problems because a substantial portion of target proteins lack known binders. Even for those targets with known binders, it is obviously advantageous to discover novel motifs in order to diversify the binder options for the purpose of affinity or property maturation. Despite the critical need, there are only few *de novo* methods that can extract structural motifs directly from protein structures. RifGen^7^ generates an extensive, human interpretable binding motifs but are too extensive to choose most promising motifs against the rest. FTmap^8^ provides most informative motif prediction, and we believe this utility can be improved in terms of prediction accuracy and application versatility by leveraging recent deep learning techniques. Other receptor-oriented methods, such as MaSIF^9^ or ScanNet^10^, provide predictions of key binding patches or residues on the target receptor, but not for the binder motifs, which differs in the utilization compared to binder-centric approaches.

In this study, we present a deep learning approach named MotifGen, for predicting most likely binding motifs solely from the input target receptor structure. More specifically, MotifGen returns a marginal probability profile of binder functional groups at every surface grid point that are human interpretable and hence can be directly incorporated into design processes. The development of a deep learning network takes advantage from recent advancement in deep learning techniques for protein structures^11,12^. With these deep learning architectures and sufficient data for binding motifs, we reasoned that the network can learn the ability to robustly estimate those binding motif types from receptor structure alone. The concept of our work can be also understood as a masked language model, but in Euclidean space^13^: the receptor structure being the context and the binding motif being the masked word.

## Results

### Overview of the work

We first summarize the network, followed by a brief description on the training process. Details of each part are further described in **Methods**. The MotifGen model features a neural network architecture designed to predict binding motifs based solely on the target receptor structure, as depicted in **Fig 1a**. The workflow begins by generating grid points on the protein surface, where an SE(3)-equivariant encoder^12^ builds grid embeddings from neighboring receptor atoms. These embeddings are then decoded into the marginal probability values of various functional groups. These functional groups, described as binding motifs, include 14 *specific types* representing functional groups located at sidechain-tips and a backbone amide group (details in **Table S1**). They are categorized into six interaction property *classes*: none, proton acceptors, proton donors, both donor and acceptor, aliphatic, and aromatic, as shown in **Fig 1b**.

**Fig 1.**
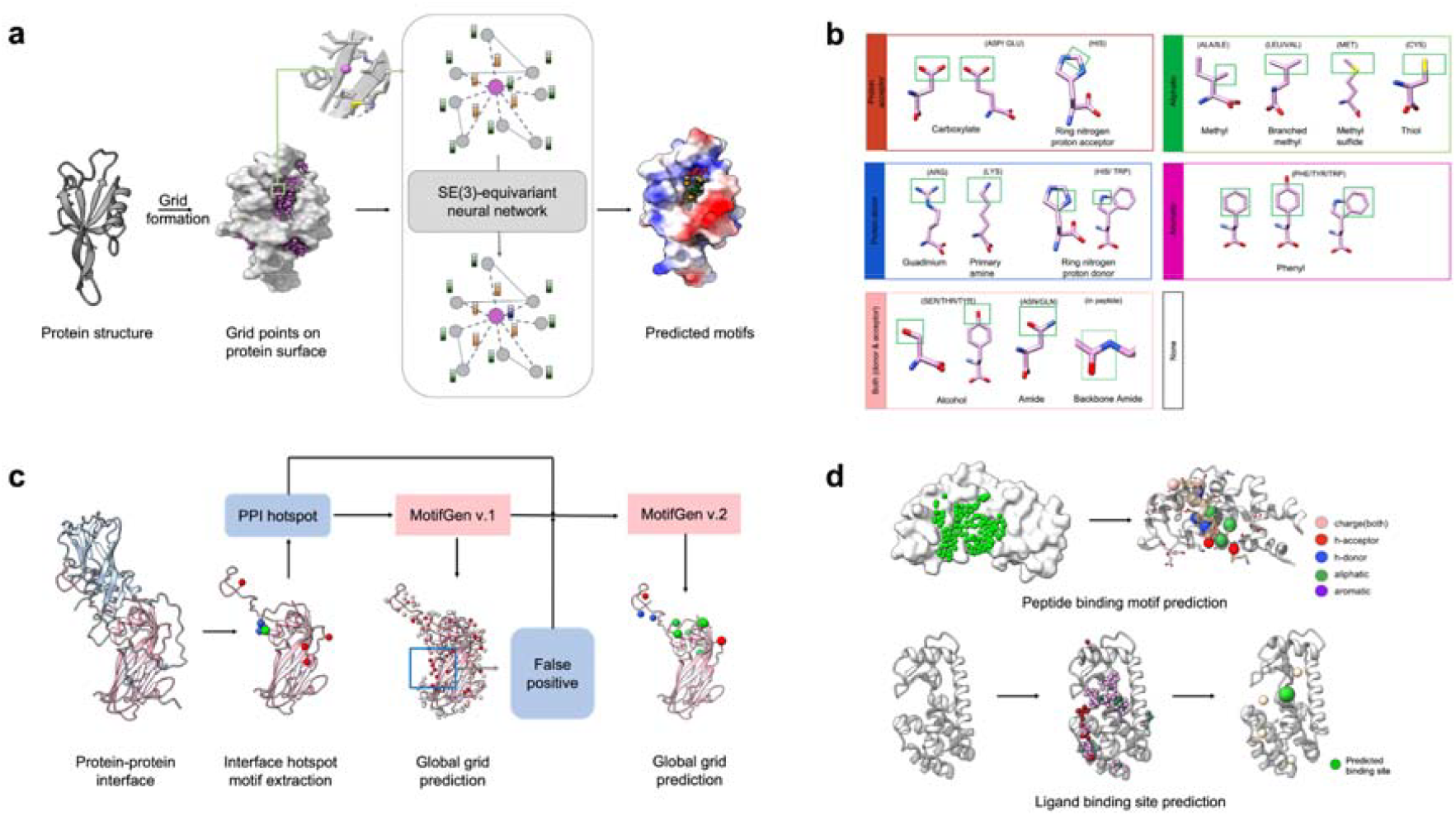
Workflow of the MotifGen model. a) Schematic representation of the MotifGen workflow. Given a protein structure, grid points are generated on the surface, where the neural network predicts marginal probability values for various functional groups. b) Definitions of binding motifs: 14 functional group *types* from canonical amino acids are represented, with motifs of the same chemical *class* grouped within the same box. Chemical functional group parts in motifs are highlighted in small green boxes. c) The training scheme of MotifGen. MotifGen is initially trained only on exact motif points extracted from protein-protein interface data. At the subsequent adversarial training stage, negative samples at other grids (i.e. non-motif site but having high probability) are added as additional training data. d) Two few-shot molecular design applications based on MotifGen profile tested in this manuscript: 1) peptide binding motif prediction and 2) ligand binding site identification.

The model is trained solely on protein-protein interface data. Total 40,543 unique clusters of protein-protein interactions are extracted from PDB biounit (Sep 28, 2020)^14^. 271,870 unique interface binding motifs are then extracted by collecting those with Rosetta energy ^15^ difference > 2.0 kcal/mol upon the chain detachment. Our rationale for using protein-protein interface data is its richness – this number is far larger than that from protein-ligand interfaces by multiple folds; note that a considerable fraction of protein-ligand complex structures are highly redundant. It is also known that interaction patterns observed in proteins can be largely generalized to protein-ligand interactions ^16,17^.

We designed the loss function to make the network favor not only specific functional group types but also those that belong to the same chemical classes sharing similar chemical properties. The loss function is designed as Lcat + Lclass, where Lcat measures the exact match to one of the 14 observed category types, while more permissive Lclass measures match to one in the same chemical class. At the first round, marginal probability of types and classes at exact motif locations are trained. After the first round training, unexpected preferences to certain types are observed at exposed grid points. We reasoned the artifact is mainly due to the lack of negative data; to penalize these false positives, adversarial negative data are collected and appended to the second round training with a ratio equal to the positive data. We observed further adversarial training beyond the second round was not necessary.

At the inference stage, the trained MotifGen generates a per-type probability profile at every solvent-accessible grid point on a protein surface. These profiles provide starting points for versatile practical applications through few-shot semi-supervised learning procedures, in which problem-specific small models are newly built. In this manuscript, two representative real-world applications are demonstrated: peptide design and protein-ligand binding site prediction. Because these applications are unrelated to the MotifGen training data, the performance from semi-supervised learning may reveal the richness and the transferability of learned motif profiles for many other downstream applications yet to be performed.

### MotifGen can generate selective binding motifs for peptide binders

We first benchmarked the predictive power of MotifGen for recovering binding motifs at known peptide-receptor complexes. We set up the problem as a local prediction, where the binding site is known or designated by a user, and the task is to determine a plausible pool of motif types and coordinates around the binding site. We believe this corresponds to a realistic scenario in drug discovery, because a designed peptide needs to bind at the desired site, rather than anywhere, in order to inhibit or modulate certain other interactions.

We set up a benchmark set consisting of 1,466 protein-peptide complexes from Propedia. We asked MotifGen to predict motif profiles at each surfacial non-clashing grid point within a 12Å radius sphere from the binding site (i.e. binder peptide center-of-mass). The benchmark result (**Fig S5**) showed small improvement in F1-score (a combined metric of precision and recall) over random assignment of one of six motif categories. We reasoned the moderate performance may originate from the difference in the interface characteristics between protein-protein (i.e. training data) and protein-peptide interfaces.

To improve the predictive power of the network for protein-peptide interfaces, starting from the pre-trained weights and the original architecture, a fine-tuning is performed on a dataset solely consisting of 1,783 protein-peptide complexes. The details of fine-tuning strategy and dataset are provided in **Methods**. The fine-tuned model resulted in further improved recovery of known binding motifs (**Fig 2a**). The model exhibited improvements across all motifs, particularly in acceptor and aliphatic categories, achieving F1 scores over 0.3 for the 1st quartile predictions. Focusing on the native-motif recovery (recall, **Fig 2b**), at least half of the actual motifs were recovered for 90% of targets. This performance demonstrates that MotifGen reliably recovers key motifs across a wide range of targets, offering valuable insights into peptide-receptor interactions. From this point, results regarding peptides will be presented using these fine-tuned weights.

**Fig 2.**
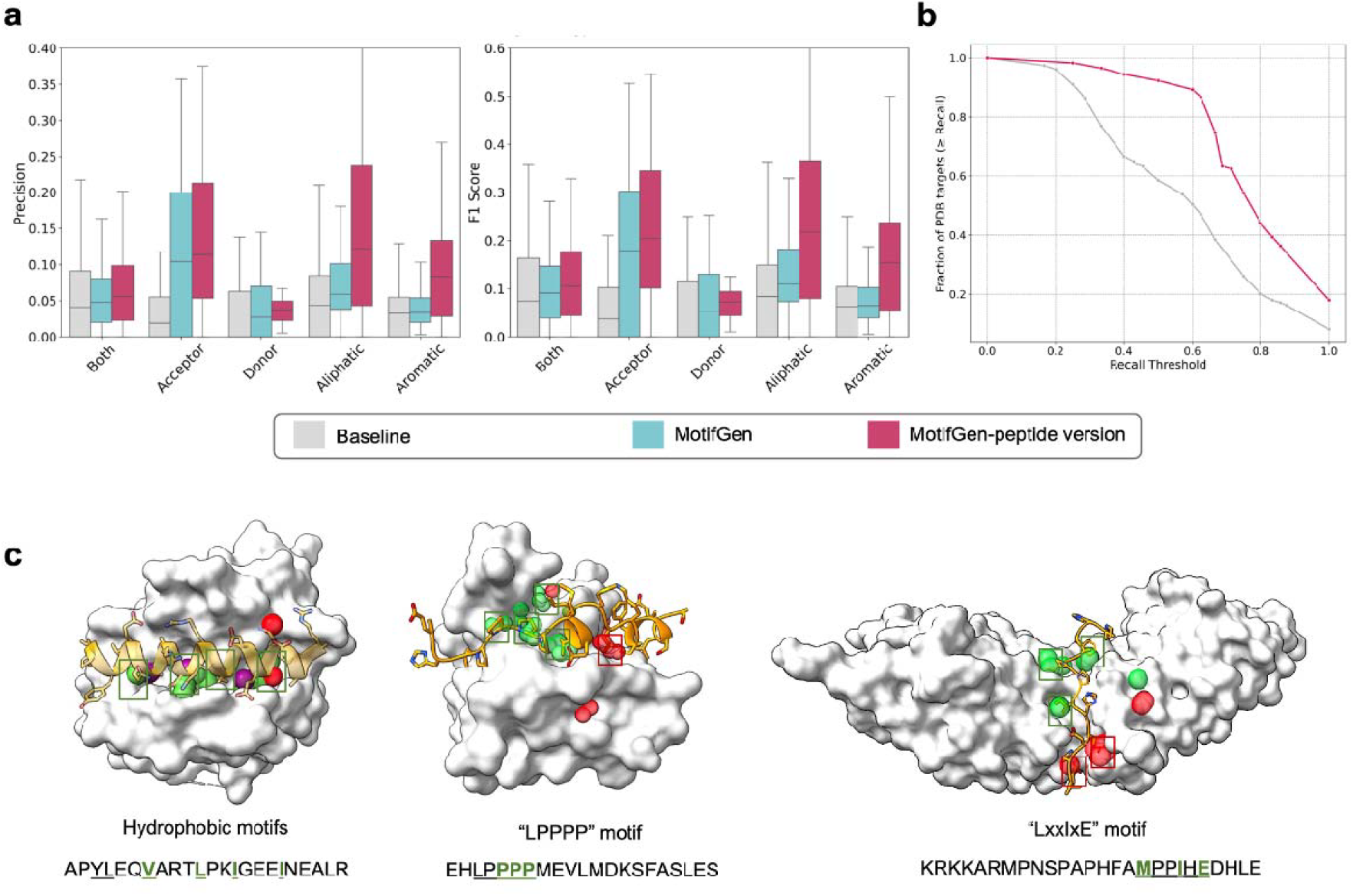
MotifGen performance on protein-peptide motif prediction. a) The motif prediction performance for a representative test set, comparing three model types: the baseline (random assignment), the model before fine-tuning using protein-peptide complexes, and the model after fine-tuning. Precision and F1 scores are collected for each interface and their statistics across the whole dataset is shown. Precision measures how accurately predictions are made within a 2.0 Å proximity to actual motifs, while the F1 score provides a balanced view of both precision and recall. b) The cumulative distribution function of recall values across the test set PDBs. The plot tracks the fraction of PDB targets that achieve at least a given recall level on the X-axis. PDBs with at least three motifs were included in this analysis. c) Examples of MotifGen predictions on three complexes in which ligands are designed peptides. (left) a peptide activator dM2 on the BAK receptor, involved in apoptosis by inducing mitochondrial outer membrane permeabilization (MOMP) and promoting cytochrome c release^18^. Key motifs are highlighted hydrophobic residues. (middle) a peptide inhibitor PCARE with the ENAH receptor in the Ena/VASP family regulating actin assembly. Key motifs involved in this binding is the LPPPP motif.^19^ (right) B56 peptide with a serine phosphatase involved in regulating phosphorylation in cellular signaling. B56 interacts with target substrates through the highlighted LXXIXE motif ^20^.

To view the predictive power in a more comprehensive way, we brought three complexes where designed peptides are bound along with the annotated binding motifs by the authors. As shown in **Fig 2c**, MotifGen accurately predicted these known motifs, aligning well with annotated motifs. These results suggest MotifGen can offer strong clues at the initial stages of peptide designs.

### MotifGen integrated with structure prediction network considerably improves peptide binder discrimination

We then assessed MotifGen’s ability for designing peptide binders given a receptor structure. There are two major tasks regarding a typical peptide design process; the first is a sequence determination task given a peptide backbone scaffold, and the second is selecting the most promising designed sequences. Here we focus on the second selection task, which is a very crucial tool for design filtering and validation.

To this end, we sought to develop an estimator that can correctly rank peptide binders and non-binders, by integrating MotifGen predictions with confidence metrics resulting from AlphaFold2^11^ (AF2). Although AF2 has become the state-of-the-art tool for modeling and validating protein-peptide complexes ^11,21^, its confidence metrics such as interface-pAE (i-pAE) or p-lDDT have been questioned for ranking design candidates^22^. We hypothesized the chemical characteristics measured by MotifGen at receptor interfaces can serve as an orthogonal metric to the AF2 confidence metrics.

To test this hypothesis, we developed a scoring network *MotifPepScore*, which returns a combined confidence score for a protein-peptide complex structure. The model takes two components as inputs: 1) AF2 predicted structure and its residue-wise confidence scores (pLDDT and PAE) and 2) *MotifGen agreement score*. The MotifGen agreement score is calculated by summing the structural overlap between the MotifGen profile and corresponding side-chain tips (**Fig 1b**) at the binder peptide model structure, measured by a Gaussian overlap function with a σ of 1 Å. These two scores are processed by a light graph self-attention network consisting of 44,509 parameters (model architecture in **Fig S2**) to return the final score.

We compared MotifPepScore against three other scores, including AF2 i-pAE, motif agreement score alone, and a MotifPepScore trained without the motif agreement term. A set of peptide binders and non-binders are brought from the PRM database^23^, containing 1,898 experimentally validated binder sequences across 18 receptor domains. Non-binder sequences are chosen at a 1:10 ratio relative to binders. In **Fig 3a**, performances of 4 scores are provided using the area-under-curve (AUC) of Boltzmann enhanced discrimination of the receiver-operator characteristics (BEDROC)^24^ to highlight their early false-rate discrimination. While both i-pAE and the MotifGen agreement score perform similarly, MotifPepScore shows a significantly improved performance in ranking peptide binders. Notably, the averaged BEDROC values across all domains reveal that MotifPepScore consistently outperforms other metrics, indicating the combination captures complementary information that neither individual metric can provide by itself.

**Fig 3.**
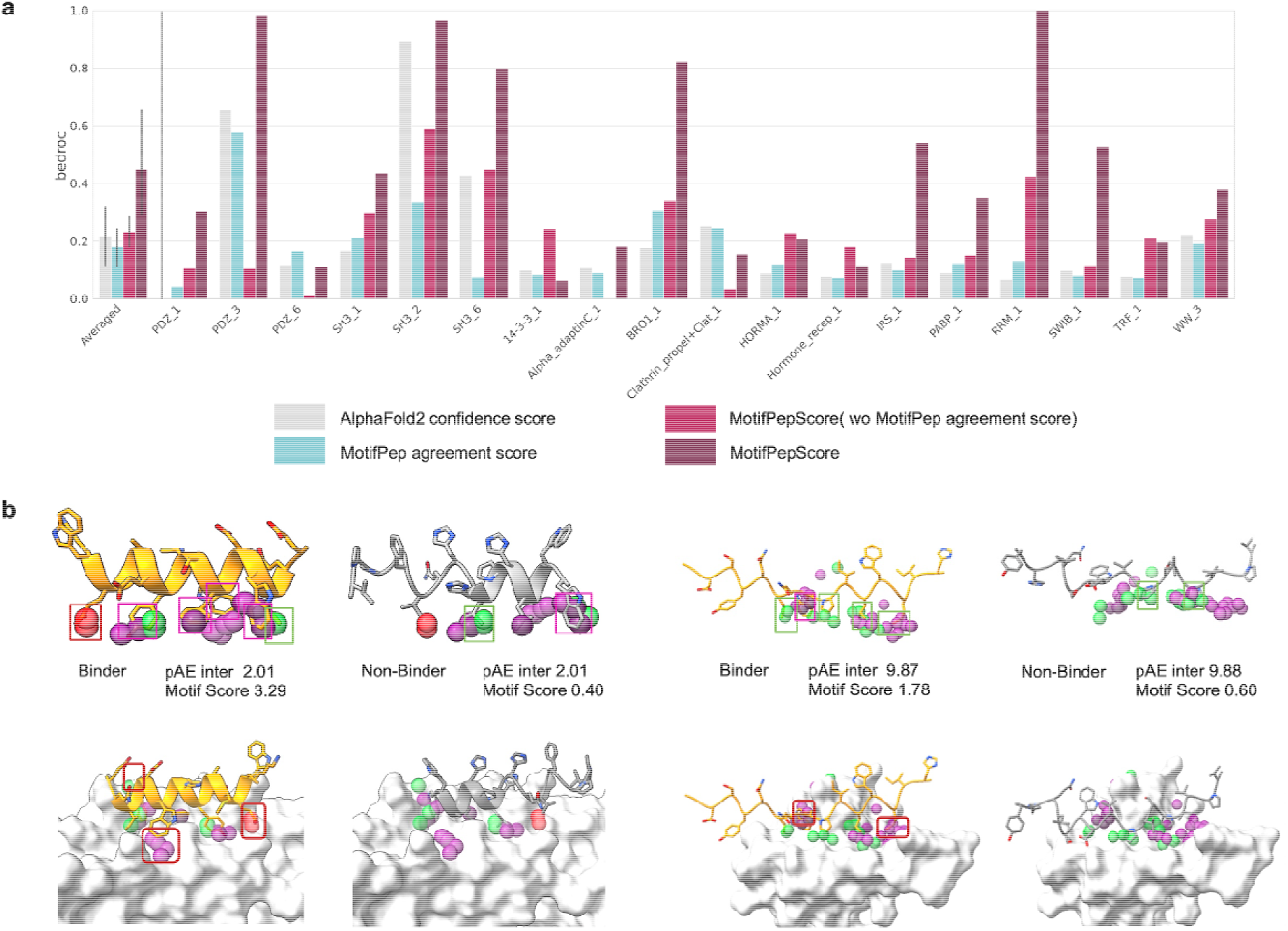
Application of MotifGen on peptide binder discrimination combined with structure prediction networks. a) Discriminative power measured by Boltzmann enhanced discrimination of the ROC curve (BEDROC). Four methods are compared on their discriminative power: i) AF2 (i-pAE), ii) MotifGen agreement score, iii) MotifPepScore without agreement score, and iv) the full MotifPepScore including the agreement score. Details of the metrics are described in the main text. b) Representative examples when addition of MotifPep agreement score helps rescue AF2-alone discrimination. Selected binders and non-binders to (left) the Bro1 domain and (right) the SWIB domain. Complex structures are modeled using AlphaFold2.

In **Fig 3b**, several examples are highlighted for which the predicted motifs work for the discrimination but typical AF2 metrics fail. The first example is the Bro1 domain, found in proteins that are involved in essential processes such as membrane fission during HIV budding, cytokinesis, and multi-vesicular body formation^25^. MotifGen predicted several key aliphatic, aromatic, and H-acceptor motifs within the Bro1 domain pocket. Examining representative binders and nonbinders, the binder peptide had all motifs with a high motif-agreement of 3.29, while the non-binder lacked some of these, resulting in a significantly lower motif-agreement of 0.40. In contrast, AF2 prediction was non-discriminative, estimating both equally stable (i-pAE equally 2.01). The second example, SWIB domain plays a crucial role in regulating transcription in the SWI/SNF complexes^26^. The MotifGen identified aliphatic and aromatic motifs, resulting motif-agreement of a selected binder by 1.78 compared to 0.60 for a non-binder, while AF2 equally predicts both weak-or non-binders (i-pAE of 9.87, 9.88 for the binder and non-binder, respectively).

We explain the improved discriminative power of MotifPepScore based on the complementarity between two components, AF2 and motif-agreement. AF2 metrics provide “global preference” based on sequence similarity and structural compatibility, but because of this, they frequently fail to capture local mutational effects ^27^. On the other hand, MotifGen focuses more on “local preference” identifying critical interaction motifs, while being weak at capturing structural compatibility. This complementary allows the combined MotifPepScore to generally perform well while also capturing single mutational effects to binding affinity.

### Accurate and generalizable prediction of small-molecule binding sites

The next application is protein-ligand binding site prediction. Here we refer to ligands as small molecules or peptides consisting at most 50 heavy atoms. Due to their small size, ligands prefer binding at pocket-shaped sites maximizing their contact area to receptors. The major question regarding our training data is, whether the set of functional groups present in 20 standard amino acids are broad and general enough to represent diverse chemistry spanned by chemical compounds. We assumed essential interactions occurring between receptors and ligands are almost identical to those found in protein-protein interactions (e.g. pi-pi stacking, hydrogen bonding, vdW) and that rest missing interaction types (e.g. halogen interactions) may have minor contributions in a large fraction of complexes^16,28^.

Given this assumption, a simple ML model named “MotifSite’’ predicting ligand binding sites is built upon the processed MotifGen profile using the original weights. First, MotifGen profiles are *globally* calculated through all solvent accessible grid points around the entire surface of a protein. Those grid points are then collected at which the probability is larger than pre-defined criteria for at least one motif type. Clustered points using DBSCAN ^29^ are ranked by a RandomForest classifier^30^ by processing per-cluster features, including maximum protrusion^41^ of the cluster, cluster size, mean and maximum marginal probability values, and number counts of each of 6 motif classes. This model is trained on a subset of PDBbind 2019^31^ refine set containing 861 complexes whose receptor sequence similarity is less than 40% to any protein in our test data.

Site prediction result is shown in **Fig 4a** with a few representative examples in **Fig 4b**. MotifSite slightly outperforms the most popular binding site prediction tool P2rank ^33^, which can be also categorized as a principle-based ML approach, giving a success rate of 83.6% (P2rank 80.0%) on COACH 420 dataset^34^, using DCA metric with a distance criteria of 4 Å. Show in **Fig 4a**, further analysis reveals the relative strength of each method; while P2rank generally performs better for nucleotides or nucleosides, MotifSite outperforms for drug-like molecules of which the number of heavy atoms are less than < 30 and ClogP range within - 4∼5. There are recent deep-learning methods specialized for binding site prediction^35,36,37^ reported to be superior to P2rank. However, because of their heavily supervised-training or evolutionary information-driven (e.g. protein language models) nature, it becomes questionable how these methods would work on predicting sites for novel targets or on allosteric sites of known targets.

**Fig 4.**
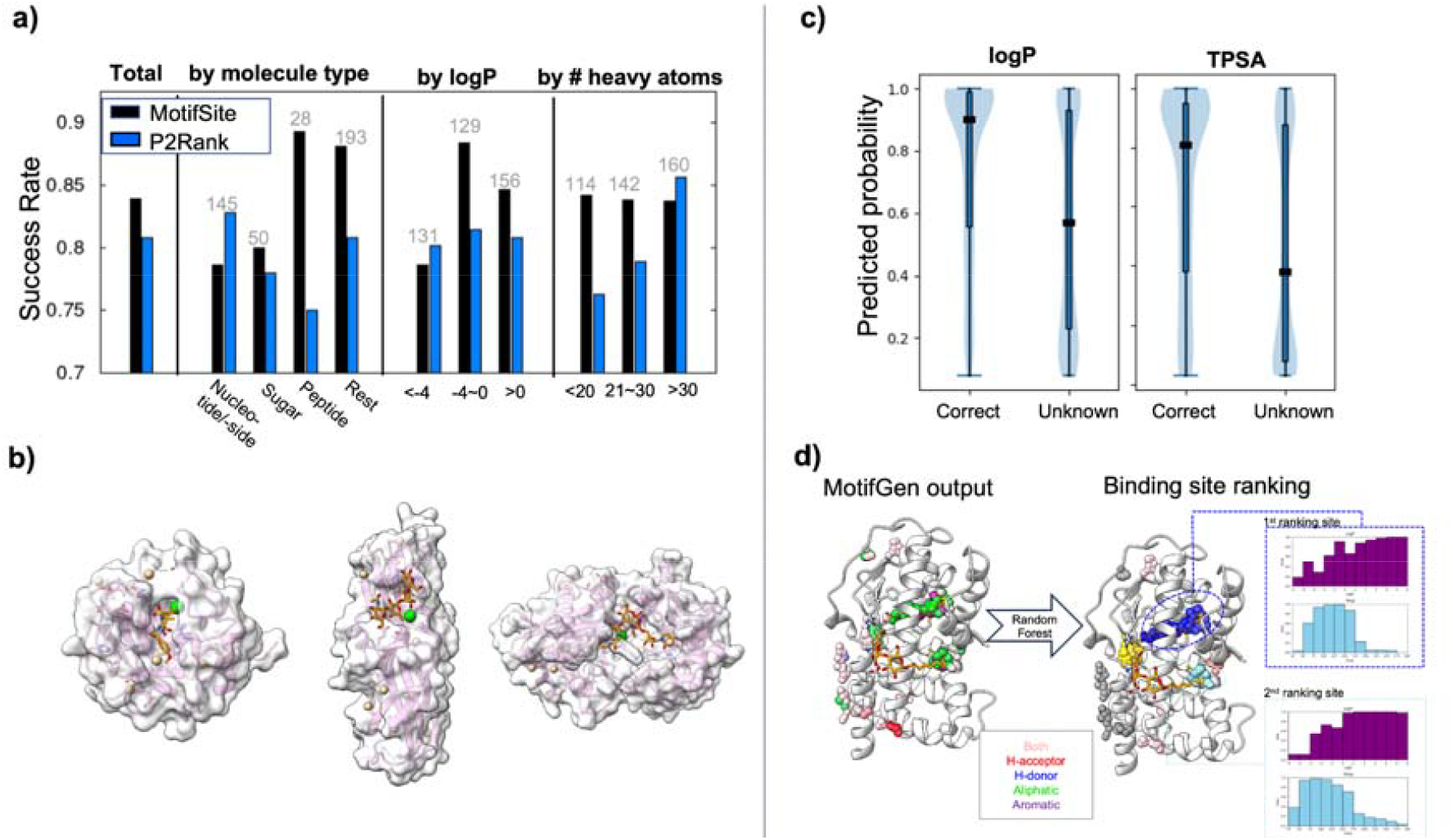
Application of MotifGen to binding site prediction. a) Binding site prediction results. Overall success rate shown on the left, with decomposed results by molecule types, by logP, and by number of heavy atoms on the right side. Results are compared over P2rank. Success is defined by DCA (distance from predicted site to any ligand atom) < 4.0 Å ^32^. **b)** Examples of binding site predictions; top-1 predicted sites are shown in green sphere, while other predicted sites are indicated in tan. The displayed proteins represent different molecule types: a drug-like ligand (PDB: 1BQO), a sugar molecule (PDB: 1J8R), and a peptide (PDB: 1P4N), from left to right, respectively. **c)** Prediction values for two binder molecular properties, logP and TPSA. “Correct” consists of value bins for ligands that are known to bind at the site, while “Unknown” bins correspond to the rest. **d)** An example of putative binder property predictions. For each site, allowed molecular properties are proposed as a range. An example is shown for PDB 1EXA.

Besides putative binding sites, predicting chemical properties of drug molecules at the target site is also of general interest in structure-based drug discovery studies.^38^ Since MotifGen not only offers most probable sites but also probable functional groups at those locations, it could be naturally extended to predicting important drug properties targeting the pocket.

Again, two RandomForest regressor models are trained upon the MotifGen profile using the same training dataset. Each model predicts possible partition coefficients (logP) and topological surface area (TPSA) values of drugs, which are considered as most essential metrics gauging the oral viability and drug absorption rates^39,40^. Regressor models predict probability values at binned ranges: for logP, from −5 to 6 with a bin size of 1; for TPSA, from 25 to 300 with a bin size of 25.

In **Fig 4c**, the range of predicted probabilities are shown for those two metrics. As shown in the figure, bins that correspond to actually observed from existing complex structures have higher probability than that for unknown values. A prediction example is shown in **Fig 4d**, for which MotifSite recommends a highly polar compound at the orthosteric site where a nucleotide binds, but a slightly hydrophobic one at an allosteric site. Using these per-site molecular property distributions, a designer can target sites where binders possess more drug-like properties, or target ligand types given a designated site.

## Discussion

In many real practice drug discovery, computational predictions should be made on unbound apo structures. To check the robustness of MotifGen, we brought all available apo structures of the identical proteins in test data. For the peptide motif prediction, using apo structures resulted in F1 scores comparable to that on peptide-bound structures (**Fig S3**). For the ligand binding site prediction, although slight decrease in prediction accuracy was observed (76%, **Table S7**), failure cases were those receptors for which large backbone or domain-level conformational changes were involved upon binding, and the rest remained consistent. These two results support MotifGen and its downstream applications are robust and generally insensitive to side-chain level changes.

Next analysis is the ablation study of MotifGen components to the prediction performance (**Fig S4**). Components considered here are as follows in the training stage: 1) the usage of classed-classification loss (Lclass), 2) the usage of adversarial negative samples, 3) the number of layers, and 4) the usage of weighted sampling inverse-proportional to the number of motif types in training data. By training the network excluding each component (reduced layers from 10 to 6 for the case 3), we find all of the factors contribute to the discrimination power, yet both motif type and class prediction performances remained robust within 10% from the optimum. Also, class prediction was reasonable even without the classed-classification loss, meaning that the network was already able to implicitly learn chemical relationships between motif types. Of the tested options, the most important part was the negative data, largely reducing the F1-score of binary prediction (i.e. whether a motif exists or not), consistent with our observation at the training stage.

In this work, we presented MotifGen and its applications to peptide sequence screening and binding site prediction. MotifGen is by philosophy similar to other deep-learning networks studying biomolecular interactions. However, unlike other receptor-centric methods, MotifGen is binder-centric and returns explicit prediction of putative binder properties, which can be readily incorporated to binder design related studies. With its demonstrated applicability to ligand binding site prediction and peptide screening, future efforts will focus on tackling more challenging tasks such as ligand virtual screening and on experimental validation of outcome designs.

## Methods

### Model architecture

#### Model overview

The network consists of 10 SE3-transformer layers^12^ followed by multiple linear layers converting virtual node’s embedding into prediction values aforementioned (**Fig S1**). For each category type, 4 linear layers are allocated: i) motif category block, ii) motif category weight block, iii) orientation weight layer, and iv) backbone displacement block. The output from the former two layers are multiplied together to return per-category marginal probability. The last two predictions were initially added in order to predict motif orientations but unused in this study.

#### Input features

Given a motif point of interest, a set of receptor heavy atoms within 12 Å to the motif point are collected and represented as graph nodes. A virtual node is allocated for the motif point. Any pair of nodes within 6 Å apart are connected by an edge. Input node features and edge features are summarized in **Table S2**. Node features include 1) 1-hot encoded amino acid type, 2) Rosetta generic atomic type^41^, 3) solvent accessibility, 4) atomic charge, and 5) distance to the virtual node. Solvent accessibility (SA) of an atom is assigned in a range [0.0,1.0] and is approximated by using the number of Cb atoms around the atom within 12 Å (N);

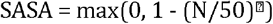

Atomic charge values are brought from the Rosetta energy parameters^42^. Distance is normalized in a range [0.0,1.0] by applying a sigmoid function:

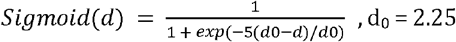

Three egde features are 1) 1-hot encoded distance (or “distogram”), 2) whether a chemical bond is formed between the atomic pair, and 3) displacement vector. Distance bins range from 0.5 to 5.0 Å with the binsize of 0.5 Å.

#### Loss function

The net loss function consists of 5 terms:

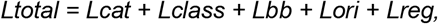

where Lcat and Lclass are loss functions for motif type and class classifications, respectively, and are evaluated as sum of binary cross entropy for each motif type or class. The rationale for using such loss function for a multiple category classification problem (instead of categorical cross entropy) is because what the network predicts is “marginal probability” rather than “joint probability”, sum of which may not necessarily sum up to 1.0 (*i*.*e*. multiple solutions are valid). In the same regard, Lclass equally rewards multiple solutions correctly that are mutually allowed based on chemical similarity (same group in **Table S1**). Lbb and Lori are losses that evaluate vector predictions, but its impact is not analyzed in this study: Lbb the backbone displacement vector from the motif centric atom, and Lori the orientation prediction (Details in **Supplementary Methods**)

### Dataset preparation

#### Collection of unique interfaces

The dataset for training was prepared using structural entries retrieved on RCSB Protein Data Bank (PDB), as of Sep 28, 2020. Structures were selected based on three criteria: having more than two polymer instances, a non-monomeric oligomeric state, and a resolution better than 3.0 Å. These criteria resulted in a total of 68,918 PDBs. To enhance the diversity of interface structures, all possible biological units were generated using the biological assembly information provided in the PDB (REMARK 350 lines). Each oligomeric structure was then systematically decomposed into its constituent dimers for subsequent processing.

These dimers are then clustered using sequence similarity. Sequences were clustered using CD-hit^43^ with a 40% sequence identity cutoff. Each dimer was assigned a sequence label representing its two constituent chains. The dimers with identical sequence labels were grouped into a single cluster, resulting in 20,647 unique dimer clusters. To introduce additional structural variation and improve training efficiency, each dimer cluster in the training set was further subdivided based on the relative orientation of its two subunits (chains). For this step, all structures within a dimer cluster were aligned using TM-align^44^ by superimposing one of the subunits. After alignment, the interface’s center of geometry was calculated for each dimer structure. The distances between these centers were then used as the basis for hierarchical clustering, with a 15 Å threshold to define geometry-based subclusters within each dimer cluster. During training, one structure was randomly sampled from each geometry-based subcluster in every epoch, allowing that the model was exposed to a wide variety of relative orientations within the same dimer cluster. This approach resulted in an extended set of 40,543 unique interface clusters.

#### Motif definition

Motifs in this work are defined as representative functional groups found in protein residues. They are grouped into 14 functional groups and 6 interaction patterns based on their chemical properties. The motif sites were collected from the hotspot residues in the protein-protein interfaces, where the hotspots are detected by using Rosetta energy change (> 2kcal/mol) upon the protein chain detachment. Total 271,870 points are collected from the dataset. Descriptions of the motif types and per-type counts in the training data are reported in **Supplementary Table S1**.

### Model training

#### Training details

The network was trained using Adam optimizer at a learning rate of 1.0e-4. Four NVIDIA A5000 GPU cards are used via torch dataparallel interface for training 217 epochs, which took about 3 days. Dropouts are applied to the node embedding at the beginning and the end of the encoder module with a rate of 0.2. In addition to dropout layers, receptor coordinates and the virtual grid point are randomly perturbed by maximum 0.2 and 0.5 Å from the input during training.

#### Augmenting negative data points

To prepare negative data for the adversarial training round, grid points are sampled not only at the interface hotspots but also at the entire protein surfaces (see below at the grid point selection section) for the training set. Marginal probability values inferenced using the first round model are evaluated, then high probability bins that are also at physically unlikely positions are collected. Physically unlikely points were either grids that may clash with protein atoms, measured by vdW radius, or grids that are highly exposed to solvent, measured following^45^. Of all negative data, 284,255 points, which approximately equals the number of positive data, were randomly chosen for the adversarial training.

#### Data point upsampling

Because the number of data points in the training set is not uniform, relatively rare motif types are upsampled by factors shown in **Table S1**, which matches the inverse of frequency in the dataset.

### Grid point selection

Grids are picked in a different manner at the training stage and for the inference. At the training stage, grid coordinates for the positive data (i.e. binding motif atoms) are brought exactly at the centric atom coordinate from PDB. For the negative data, grids are evenly distributed through a 3-D voxel with a bin size of 1.5 Å. At the inference stage, to predict on most likely positions for motifs, grid points are distributed at the solvent accessible points ^46^, using the probe radius of 1.65 Å and Nsample-per-grid of 50.

### Fine-tuning weights for applications to peptide study

#### Dataset

Protein-peptide interactions are extracted from Propedia 2.3 dataset, which comprises 1,466 representative protein-peptide interfaces. Since these 1,466 complexes were derived from clustered results, the dataset was randomly split into training, validation, and test sets with a 7:1:2 ratio. Validation set was also used to determine optimal threshold for each motif maximizing the F1 score of motif prediction.

#### Fine-tuning

The fine-tuning process was performed using the Adam optimizer with a decreased learning rate of 1.0e-5. The fine-tuning was run for 13 epochs starting from 217-th epoch parameters in the original model. Only the final two SE(3)-transformer layers (of the total 10 layers), the motif category block, and the motif category weight block were updated, while all the rest parameters were frozen. Adversarial negative data were introduced at a 5-fold ratio to positive data (originally 1:1 ratio) from the beginning of the tuning. The majority of the tuning process followed the original process with a few differences below. The loss function was modified to double the weight to the *Lclass* loss to prioritize chemically similar motif groups. Grid points were restricted to the points that are within a 12 Å radius to the peptide’s center of mass (originally entire protein), concentrating on probable interaction sites near peptide-protein interfaces.

### Training MotifPepScore for peptide sequence discrimination

#### Dataset

To train the MotifPepScore model for peptide binder and non-binder discrimination, we curated a dataset from the PRM database^23^, comprising 1,783 experimentally validated peptide binders across 18 receptor domains. Non-binder sequences were brought from binders to other receptors in the database with sequence similarity to the target receptor less than 40%. Binder-to-nonbinder ratio are fit to 1:10. The domain groupings, domain numbers, PRM IDs, number of binders, and fold assignments used for cross-validation are listed in **Table S3**.

#### Motif agreement score

The score evaluates the structural overlap between the MotifGen predictions and the motifs from AF2 prediction model structures. This score increases as the predicted peptide motif site at the AF2 model get closer to the predicted motif locations.The total score is a sum of individual motif class scores (e.g., proton acceptor, donor, aliphatic, aromatic), which is measured by Gaussian overlap to all grid points weighted by their motif probability on the category (p_i,c_):

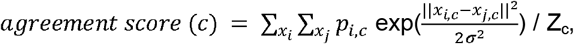

where c represents category type, x the coordinates; i grid points with motif probabilities; j atomic points in the AF2 model. The normalization factor Z, which counts the number of grid points with p_i,c_> p_threshold,c_ (per-category probability threshold reported in **Supplementary Table S6**) is introduced to measure the strength of predictions of the category.

#### The MotifPepScore network

The network integrates structural aspects extracted from AF2 and the pre-computed motif profiles from MotifGen (using peptide-fine-tuned weights) to return the likelihood of the peptide model structure as a binder. The network first constructs a graph by taking AF2 model structure, in which each peptide residue is connected to the 10 nearest receptor residues and to all other peptide residues. Input features used in the network are further described in **Table S4**. The model has a simple neural network architecture consisting of 44,509 parameters, in which a series of layers are applied as following: the first graph attention updates peptide node features followed by a self-attention layer to aggregate features, then subsequently passed through the MLP classifier to predict peptide binding (**Fig S2**).

The model was trained using a cross-entropy loss, with performance evaluation using 5-fold cross-validation due to large discrepancy between dataset for different receptors. In the cross-validation strategy, five folds were distributed to training, validation, test, with a ratio of 3:1:1. This approach was repeated for all five folds, ensuring that every fold served as the test set exactly once.

### Binding site detection module

#### Overall procedure

First, the grid points are filtered where MotifGen probability values are above the predefined thresholds: 0.25 for ASP and backbone; 0.2 for AMD, PHE, PHO, and PH2; 0.15 for LYS; rest types are unused. These grid points are clustered using DBSCAN ^29^ with epsilon=2.4 and min_samples=5. Per-cluster features are extracted, which contains chemical and geometrical properties of the cluster (full list in **Table S5**).

Random forest classifiers were separately trained on these features to predict binding sites and binder properties, respectively. For the binding site classifier, per-cluster features were taken as input, and whether each cluster center-of-mass locates within < 4.0 Å to any ligand atom (DCA, distance from site center to atom) was trained against the dataset described below. To label multiple binding sites in the training data correctly, we searched against bioLip database (until Dec 1st, 2022)^47^, and added labels if any drug-like molecules with DCA < 4.0 were found at the site.

Similar procedures were carried out for training binding property predictors. First, “known” ligand properties were collected from the bioLip database from those ligands sharing same binding sites. For those ligands, TPSA (topological polar surface area) and ClogP (calculated partition coefficient) were calculated using obabel (v 3.1.0). Binned labels were used; from 25 to 300 with the bin size of 25 for TPSA labels; from −4.0 to 5.0 with the binsize of 1.0 for ClogP. Bins at known properties or their neighbors were marked as 1.0; otherwise 0.0 was assigned. Using these as labels, marginal probability values at every bins were trained.

For all Random Forest models, the parameter set of random_state=0, depth=10, n_estimators=500 were used. We tested several different parameter combinations, but their difference was only 1∼2%, meaning that the training was robust regardless of the model parameter setup. The model was also trained on different sizes of training data from 100 to full (861) receptors, but the model trained on the smallest set performed reasonably well, with only 3% decrease in binding site prediction.

#### Dataset

COACH420 set^34^ was taken as the test set for direct comparison to published methods. Training data was brought from the refined set of PDBbind set v2019^31^. All targets whose receptor similarity is higher than 40% to any test targets were excluded, leaving 861 (of total 4852) proteins-ligand complexes for the training data. The apo structures of the test set were brought from APObind^48^). Of 420 test set receptors, 95 were taken for the test which have their corresponding apo structures in APObind.

## Notes

### Competing Interest Statement

The authors have declared no competing interest.

