## Supplementary Material for "Deep learning molecular interaction motifs from Receptor structure alone"

Authors: Seeun Kim[1], Simaek Oh[1], Hyeonuk Woo[1], Jiho Sim[1], Chaok Seok[1]*, Hahnbeom Park[2,3]*
[1] Department of Chemistry, Seoul National University, Seoul 08826, Republic of Korea
[2] Biomedical Research Division, Korea Institute of Science and Technology, Seoul 02792, Republic of Korea
[3] KIST-SKKU Brain Research Center, SKKU Institute for Convergence, Sungyunkwan University, Suwon 16419, Republic of Korea.

#### Contents

**Supplementary Figures**
**Fig S1.** MotifGen architecture
**Fig S2.** MotifPepScore network architecture
**Fig S3.** Comparison of MotifGen Prediction Accuracy on Modeled vs. Holo Receptors.
**Fig S4.** Ablation study of MotifGen

**Supplementary Tables**
**Table S1**. Definitions of 14 binding motifs
**Table S2.** Input features used in the MotifGen network
**Table S3.** The PRM dataset processed for training MotifPepDesignScore.
**Table S4.** Input features used in the MotifPepScore network
**Table S5.** Cluster input features used in the MotifSite classifiers
**Table S6.** Per-motif category probability threshold for calculating the motif agreement score
**Table S7.** Apo state binding site prediction results

**Supplementary Methods.**
1) Definitions of auxiliary losses

### Supplementary Figures

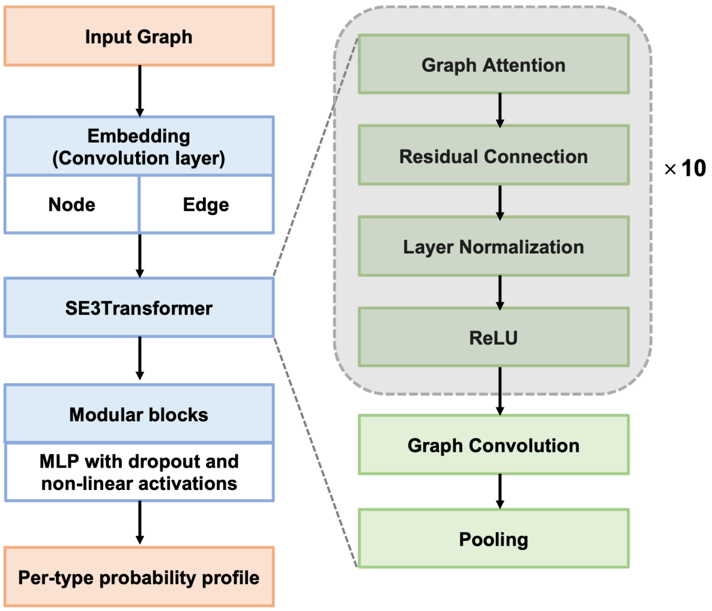


**Fig S1.** MotifGen architecture.

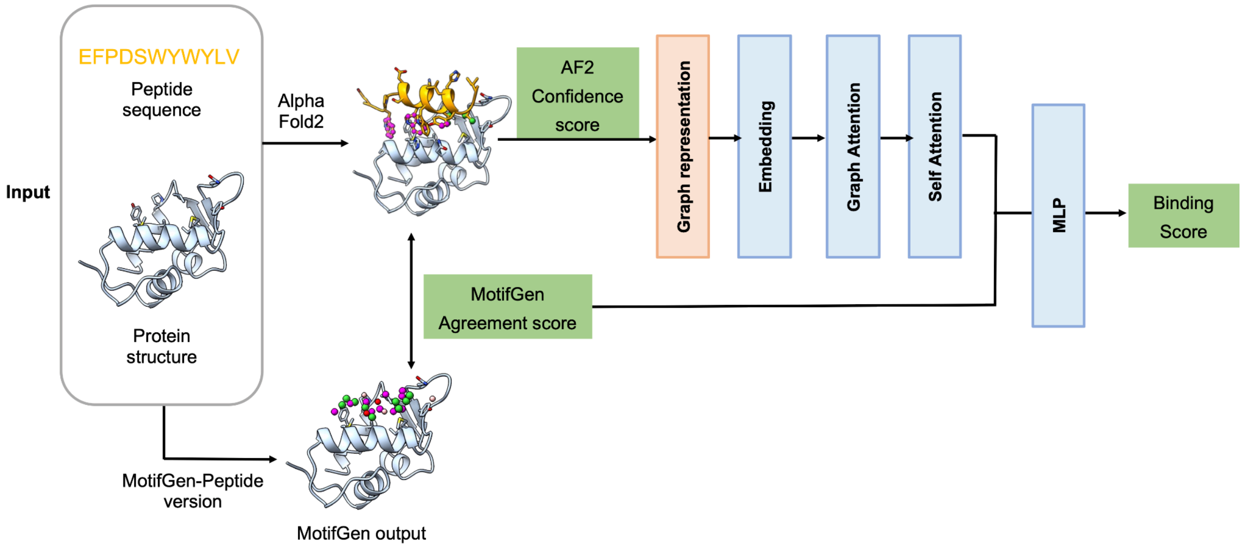


**Fig S2.** MotifPepScore network architecture.

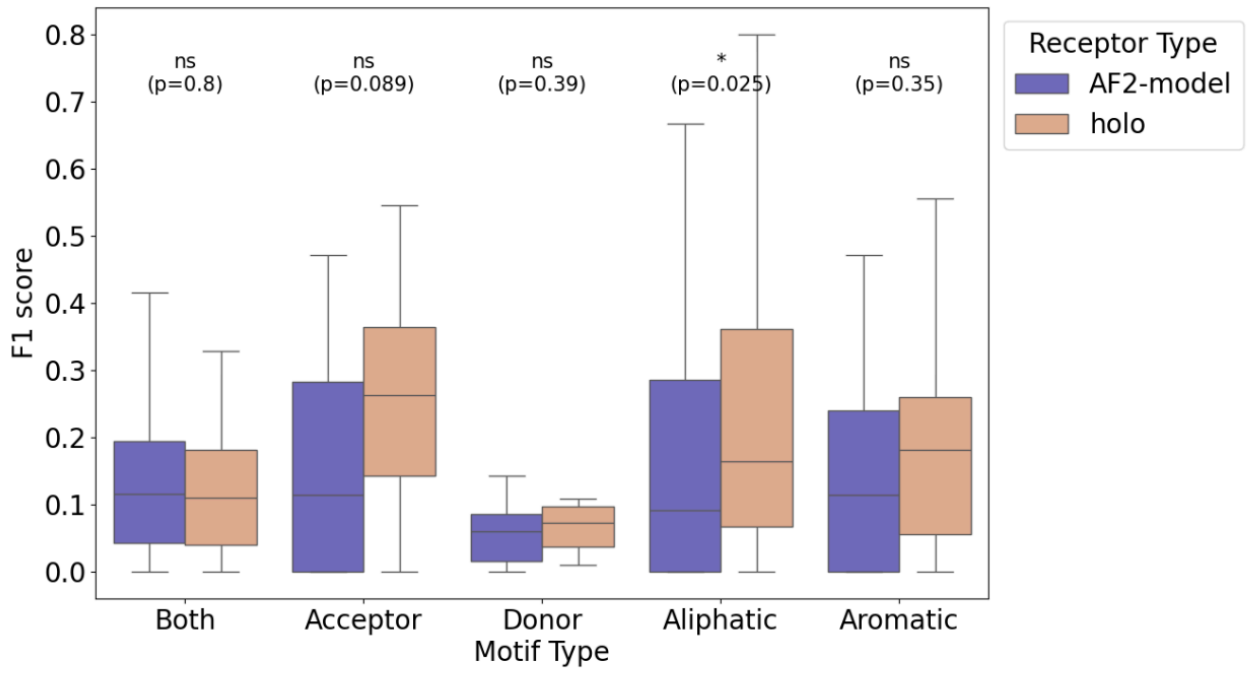


**Fig S3. Comparison of MotifGen Prediction Accuracy on Modeled vs. Holo Receptors.** Evaluation of MotifGen accuracy on modeled receptors (mean binding site pLDDT > 95) generated using AlphaFold2, compared to holo receptors. Differences in performance were assessed using the Wilcoxon Rank-Sum Test, with results annotated as text on the figure (ns for $p>0.05$, * for $0.01< p \leq 0.050$).

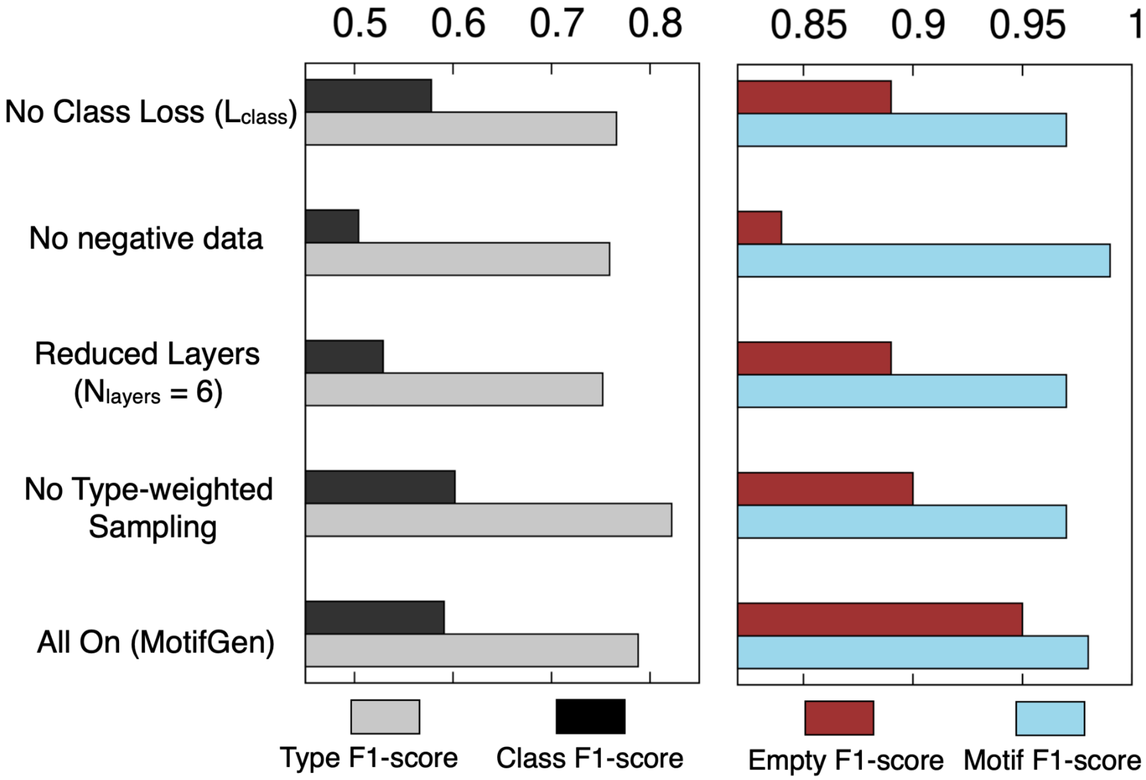


**Fig S4. Ablation study of MotifGen.** Four conditions are tested: 1) No Class-Loss, 2) No negative data, 3) Reduced encoder layers (from 12 to 6), and 4) No type-weighted sampling (i.e. uniform weights across types). Left) F1-score values on type predictions (gray) and class predictions (black). Right) Binary prediction F1-score values on empty site predictions (type=0, dark-red) and site where any motif exists (type>0, cyan).

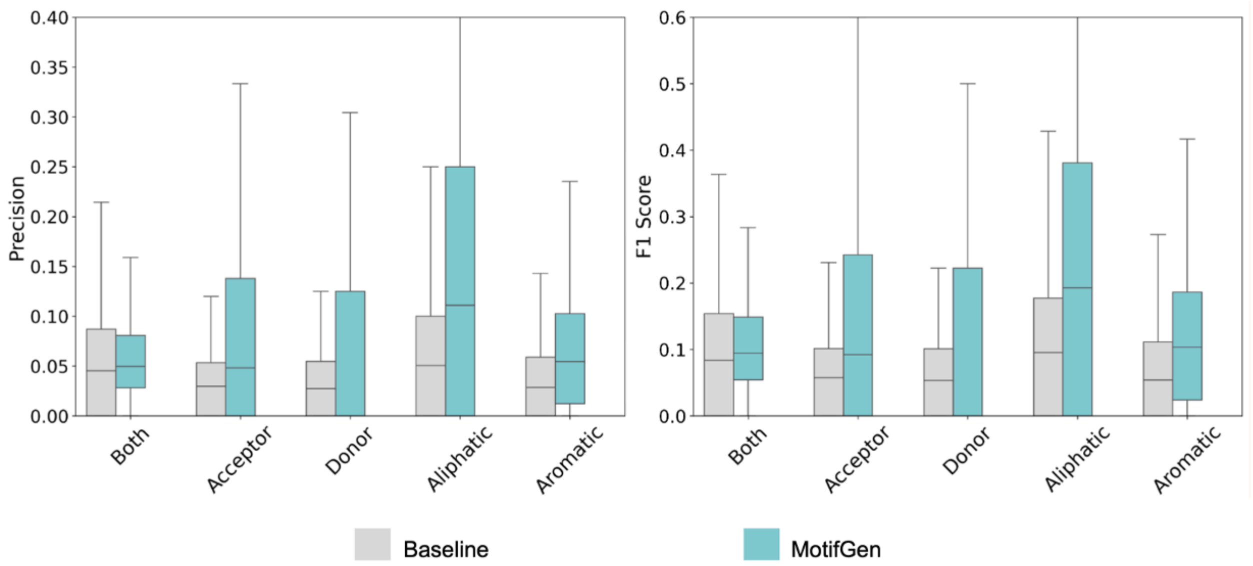


**Fig S4. Benchmarking MotifGen on Protein-Peptide Complexes from Propedia.**

#### Supplementary Tables

**Table S1. Definitions of 14 binding motifs**

| Name | Functional group | Correspondi ng amino acids | Centric atoms | Number of data points | Sampling weight | Class index |
|---|---|---|---|---|---|---|
| None | None | - | - | 284,255 | 1.0 | 0 |
| ASP | Carboxylate | ASP, GLU | C | 20,841 | 3.0 | 2 (acceptor) |
| ARG | Guadinium | ARG | C | 7,078 | 5.0 | 4 (donor) |
| LYS | Primary Amine (protonated) | LYS | N | 6,665 | 8.0 | 4 (donor) |
| OH | Alcohol | SER, THR, TYR | O | 17,030 | 4.0 | 3 (donor & acceptor) |
| BB | Amide (in peptide) | All | C | 25,274 | 4.0 | 3 (donor & acceptor) |
| AMIDE | Amide | ASN, GLN | C | 13,714 | 6.0 | 3 (donor & acceptor) |
| NHIS | Ring nitrogen proton acceptor | HIS | N | 15,585 | 6.0 | 2 (acceptor) |
| NTRP | Ring nitrogen proton donor | TRP, HIS | N | 8,615 | 6.0 | 4 (donor) |
| PHE | Phenyl | PHE, TYR, TRP | CG | 35,915 | 2.0 | 5 (apolar) |
| SH | Thiol | CYS | S | 0 (unused) | 1.0 | 5 (apolar) |
| CH3 | Methyl | ALA, ILE | C | 65,363 | 1.5 | 5 (apolar) |
| CH32 | Branched methyl | LEU, VAL | Central C | 75,473 | 1.5 | 5 (apolar) |
| SCH3 | Methyl sulfide | MET | S | 10,317 | 6.0 | 5 (apolar) |
| Total | | | | 556,125 | | |

**Table S2.** Input features used in the MotifGen network

| | Feature | Description |
|---|---|---|
| Node feature (Receptor heavy atoms within 12 Å of the motif point, plus a virtual node at the motif center) | Amino acid type | One-hot encoding of the residue type |
| | Rosetta generic atomic type | Atom types discretized according to Rosetta's generic scheme, then one-hot encoded |
| | Solvent accessibility | Ranges in [0,1]. Approximated by counting the number of Cβ atoms within a 12 Å neighborhood |
| | Atomic charge | Atomic charge derived from atom type and residue type |
| | Distance to the virtual node | Euclidean distance from each atom to the motif center (virtual node) |
| Edge feature (pair of nodes within 6 Å apart) | 1-hot encoded distance(distogram) | Distance bins range from 0.5 to 5.0 Å with the binsize of 0.5 Å. |
| | Chemical bond or not | Binary indicator for whether two atoms are chemically bonded or not |
| | Displacement vector | A 3D vector $\vec{x}_i - \vec{x}_j$ representing the relative position/direction from node $i$ to $j$ |

**Table S3. The PRM dataset processed for training MotifPepDesignScore.**

| domain group | domain number | PRM ID | # of binders | k-fold(for training) |
|---|---|---|---|---|
| 14-3-3 | 1 | PRM_0564 | 102 | 1 |
| Clathrin_propel+Clat | 1 | PRM_0516 | 50 | 1 |
| SH3 | 2 | PRM_0156 | 137 | 1 |
| RRM_1 | 1 | PRM_0600 | 141 | 1 |
| TRF | 1 | PRM_0589 | 116 | 2 |
| Hormone_recep | 1 | PRM_0601 | 143 | 2 |
| PDZ | 1 | PRM_0435 | 50 | 2 |
| SH3 | 6 | PRM_0239 | 122 | 3 |
| BRO1 | 1 | PRM_0592 | 146 | 3 |
| PDZ | 3 | PRM_0372 | 50 | 3 |
| IRS | 1 | PRM_0526 | 114 | 3 |
| Alpha_AdaptinC2 | 1 | PRM_0533 | 54 | 4 |
| HORMA | 1 | PRM_0522 | 138 | 4 |
| SH3 | 1 | PRM_0157 | 46 | 4 |
| SWIB | 1 | PRM_0514 | 140 | 4 |
| PABP | 1 | PRM_0588 | 26 | 5 |
| PDZ | 6 | PRM_0380 | 72 | 5 |
| WW | 3 | PRM_0070 | 136 | 5 |
| Total | | | 1,783 | |

**Table S4. Input features used in the MotifPepScore network**

| | Feature | Description |
|---|---|---|
| receptor node feature (10 nearest receptor residues for each peptide residue) | aatype | One-hot representation of the receptor residue amino acid sequence (20 + unknown 1) |
| | ESM2 embedding | output from layers of the ESM2-t33-650M model (layer33). |
| peptide node feature | aatype | One-hot representation of the peptide residue amino acid sequence (20 + unknown 1) |
| | ESM2 embedding | output from layers of the ESM2-t33-650M model (layer33). |
| | positional embeddings | sinusoidal positional encoding for each peptide residue |
| | pLDDT | pLDDT confidence score from AlphaFold2 |
| Edge feature | Distance | min distance between peptide residue and receptor residue |
| | Predicted Aligned Error | pAE confidence score from AlphaFold2 |
| | Distogram | predicted distogram from AlphaFold2 |

**Table S6. Cluster input features used in the MotifSite classifiers**

| **Feature name** | **Description** |
|---|---|
| Protrusion | Exposure to solvent following Ref X |
| Max Probability | Max value of motif probability in the cluster |
| Mean Probability | Mean value of motif probability in the cluster |
| Mean Hydrophilicity | Number of AMD, ASP, LYS, backbone motifs in the cluster, divided by total num members |
| Mean Hydrophobiciy | Number of PHE, PH2, PHO motifs in the cluster, divided by total num members |
| Mean positive charge | Number of ASP motifs in the cluster, divided by total num members |
| Mean negative charge | Number of LYS motifs in the cluster, divided by total num members |

**Table S6. Per-motif category probability threshold for calculating the motif agreement score.**

| Category | Threshold | Category | Threshold |
|---|---|---|---|
| ASP | 0.32 | NHIS | 0.9 |
| ARG | 0.05 | NTRP | 0.4 |
| LYS | 0.3 | PHE | 0.48 |
| OH | 0.1 | SH | 0.2 |
| BB | 0.1 | CH3 | 0.45 |
| AMIDE | 0.5 | CH32 | 0.35 |
| SCH3 | 0.9 | | |

**Table S7. Apo state binding site prediction results**

| PDBID | P2rank | MotifGen | MotifGen, apo | PDBID | P2rank | MotifGen | MotifGen, apo |
|---|---|---|---|---|---|---|---|
| 1a4kH | 1 | 1 | 0 | 1ohrA | 1 | 1 | 1 |
| 1a7xA | 0 | 0 | 0 | 1onhA | 1 | 1 | 1 |
| 1a8tA | 1 | 1 | 0 | 1ppcE | 1 | 1 | 1 |
| 1afkA | 1 | 0 | 0 | 1pphE | 1 | 1 | 1 |
| 1bnwA | 1 | 1 | 1 | 1qbuA | 1 | 1 | 1 |
| 1bqoB | 1 | 1 | 1 | 1qjiA | 1 | 1 | 1 |
| 1br6A | 0 | 1 | 1 | 1ql9A | 1 | 1 | 1 |
| 1bzyA | 1 | 1 | 1 | 1qtiA | 1 | 1 | 1 |
| 1cetA | 0 | 0 | 1 | 1swkA | 1 | 1 | 1 |
| 1ch8A | 1 | 1 | 1 | 1uvtH | 1 | 1 | 1 |
| 1cimA | 1 | 1 | 1 | 1vcuA | 1 | 1 | 0 |
| 1d3pB | 1 | 1 | 0 | 1x6uA | 1 | 1 | 1 |
| 1d4pB | 1 | 1 | 0 | 1xz8A | 1 | 1 | 1 |
| 1e3vA | 1 | 1 | 1 | 1ydsE | 0 | 1 | 1 |
| 1efyA | 1 | 0 | 0 | 1ydtE | 1 | 1 | 1 |
| 1ettH | 1 | 0 | 1 | 1zajA | 1 | 1 | 1 |
| 1eveA | 1 | 1 | 1 | 2chzA | 0 | 0 | 0 |
| 1ex8A | 1 | 1 | 1 | 2g97A | 1 | 1 | 0 |
| 1ezqA | 1 | 1 | 0 | 2ggaA | 1 | 1 | 1 |
| 1f0tA | 1 | 1 | 1 | 2gj5A | 1 | 1 | 0 |
| 1f7bA | 1 | 1 | 1 | 2hobA | 1 | 1 | 0 |
| 1fjsA | 1 | 1 | 0 | 2hxmA | 1 | 1 | 1 |
| 1fkgA | 1 | 1 | 1 | 2rfhA | 1 | 1 | 1 |
| 1g4oA | 1 | 1 | 1 | 2rkmA | 0 | 1 | 1 |
| 1gwmA | 1 | 1 | 0 | 2uyqA | 0 | 0 | 0 |

| | | | | | | | |
|---|---|---|---|---|---|---|---|
| 1h1pA | 1 | 1 | 1 | 2vaqA | 1 | 1 | 1 |
| 1h1sA | 1 | 1 | 1 | 2vcjA | 1 | 1 | 1 |
| 1hdqA | 1 | 1 | 1 | 2vkmA | 1 | 1 | 0 |
| 1hpvA | 1 | 0 | 1 | 2wvaA | 1 | 1 | 0 |
| 1htfA | 1 | 1 | 1 | 2x4oA | 0 | 1 | 1 |
| 1i7zA | 1 | 1 | 1 | 2zasA | 1 | 1 | 1 |
| 1i8zA | 1 | 1 | 1 | 2zgmB | 1 | 0 | 1 |
| 1if7A | 0 | 1 | 1 | 2zu3A | 1 | 1 | 1 |
| 1jq3A | 1 | 1 | 1 | 3aaqA | 1 | 1 | 1 |
| 1jsvA | 0 | 0 | 0 | 3cwkA | 1 | 1 | 1 |
| 1k1jA | 1 | 1 | 1 | 3erkA | 1 | 1 | 1 |
| 1k22H | 1 | 1 | 1 | 3gpoA | 1 | 1 | 1 |
| 1kv1A | 1 | 1 | 1 | 3hitA | 1 | 1 | 1 |
| 1kv2A | 1 | 1 | 1 | 3i6cA | 1 | 1 | 1 |
| 1l8gA | 1 | 1 | 1 | 3iesA | 1 | 1 | 1 |
| 1lqdB | 1 | 1 | 1 | 3lbzA | 0 | 1 | 1 |
| 1m48A | 0 | 0 | 0 | 3stdA | 1 | 1 | 1 |
| 1mq5A | 0 | 1 | 1 | 4stdA | 1 | 1 | 1 |
| 1mq6A | 0 | 1 | 0 | 5stdA | 1 | 1 | 1 |
| 1nhuA | 0 | 0 | 0 | 5tlnA | 1 | 1 | 1 |
| 1nhvA | 0 | 0 | 0 | 830cA | 1 | 1 | 1 |
| 1nmkA | 1 | 1 | 1 | 966cA | 1 | 1 | 1 |
| 1nz7A | 1 | 1 | 1 | Total | 79 | 82 | 72 |

**Supplementary Methods**

1) Definition of auxiliary losses

Here we define Lori for motif orienation prediction and Lbb for backbone displacement vector prediction from a motif site. Backbone displacement vectors can be quite uncertain. Therefore instead of using stiff functions (such as MSE), we took a Gaussian mixture loss (consists of 3 Gaussian functions) to allow multiple solutions:

$$\text{Lbb} = \sum_{i}^{3} \frac{1}{\sigma_i(\theta)} exp(-(\frac{v_i(\theta)-\hat{v}}{\sigma_i(\theta)})^2)$$

where $\hat{v}$ is the label displacement vector; $v_i(\theta)$, and $\sigma_i(\theta)$ are predicted vectors and sigma for i-th Gaussian function. The minimal value of $\sigma_i(\theta)$ is[41] capped to 1.0. Contribution from each Gaussian function to the total loss is proportional to their estimated confidence $1/\sigma_i$. Finally, Lreg is the regularization loss which suppresses overall weight values, which is defined as the sum of abs(parameter weights) times 1.0e-5.
